# Excretion of triacylglycerol as a matrix lipid facilitating apoplastic accumulation of a lipophilic metabolite shikonin

**DOI:** 10.1101/2021.08.04.455005

**Authors:** Kanade Tatsumi, Takuji Ichino, Natsumi Isaka, Akifumi Sugiyama, Yozo Okazaki, Yasuhiro Higashi, Masataka Kajikawa, Hideya Fukuzawa, Kiminori Toyooka, Mayuko Sato, Ikuyo Ichi, Koichiro Shimomura, Hiroyuki Ohta, Kazuki Saito, Kazufumi Yazaki

## Abstract

Plants produce a large variety of lipophilic metabolites, many of which are secreted by cells and accumulated in apoplasts. The mechanism of secretion remains largely unknown, because hydrophobic metabolites, which may form oil droplets or crystals in cytosol, inducing cell death, cannot be directly secreted by transporters. Moreover, some secondary metabolic lipids react with cytosolic components leading to their decomposition. Lipophilic metabolites should thus be solubilized by matrix lipids and compartmentalized by membrane lipids. The mechanism of lipophilic metabolite secretion was assessed using shikonin, a red naphthoquinone lipid, in *Lithospermum erythrorhizon*. Cell secretion of shikonin also involved the secretion of about 30% of triacylglycerol (TAG), composed predominantly of saturated fatty acids. Shikonin production was associated with the induction of large amounts of the membrane lipid phosphatidylcholine. Together with *in vitro* reconstitution, these findings suggest a novel role for TAG as a matrix lipid for the secretion of lipophilic metabolites.

## INTRODUCTION

Lipids are essential constituents of all living cells. Unlike the other major constituents of cells (proteins, carbohydrates, and nucleic acids), lipids are loosely defined based on their physical properties, specifically, their hydrophobicity. Lipids can be extracted from plant cells with nonpolar organic solvents such as chloroform. Structurally, this class of compound is extremely diverse, ranging from low molecular weight metabolites to polymers like cutin. Most plant lipids of low molecular weight can be roughly divided into two types: fatty acid-derived lipids and isoprenoid compounds, synthesized from fatty acids by the glycerolipid biosynthetic and isoprenoid pathways, respectively (Ohlrogge and Browse, 1995). The primary metabolites triacylglycerol (TAG) and phospholipids function to store energy and as membrane components, respectively. Fatty acid-derived metabolites may also be used in the synthesis of signaling molecules, such as jasmonic acid, a plant hormone that plays a critical role in defense reactions. Isoprenoid compounds are particularly abundant and diverse in plants, with more than 50 000 compounds identified to date. Plant isoprenoids include specialized (secondary) metabolites, which participate in interactions between plants and organisms in their environment, including insects, fungi, and bacteria (Bartley and Scolnik, 1995; McGarvey and Croteau, 1995; Pulido et al., 2012). These lipophilic isoprenoids enhance the ability of individual plants to adapt to their habitats, for example, by defending plants against other biotic and abiotic stresses and by attracting beneficial organisms like pollinators (Yazaki et al., 2017).

Unlike water-soluble metabolites that generally accumulate in vacuoles, lipophilic compounds are often transported to extracellular spaces, such as epicuticular cavities in trichomes and apoplastic spaces of oil glands, in which they accumulate (Balcke et al., 2017). For example, citrus species produce large amounts of monoterpenes and furanocoumarins, which accumulate specifically in the oil cavities of pericarps, which are apoplastic spaces surrounded by epithelial cells (Voo et al., 2012). To date, however, the molecular mechanisms underlying both the excretion and accumulation of these hydrophobic secondary metabolites remain largely unknown (Samuels et al., 2008; Tissier et al., 2017). Biochemical approaches available to analyze the secretion of lipid molecules are limited, because these molecules are secreted by only certain types of cells, such as secretory cells in glandular trichomes and epidermal cells, and obtaining sufficient numbers of these cells for biochemical analysis is difficult. The secretion of lipids from plant cells also beneficial that lipophilic metabolites remain stable in apoplastic spaces.

To analyze these fundamental biological events, we have used the shikonin production system of *Lithospermum erythrorhizon*. The roots of *L. erythrorhizon*, which produce large amounts of shikonin derivatives, have been traditionally used as a crude drug in East Asian countries. Shikonin derivatives have a variety of biological activities, including antibacterial, wound-healing, and anti-inflammatory properties and have been found to have previously undetected pharmacological activities, such as anti-topoisomerase activities (Yazaki, 2017). These red naphthoquinone derivatives, which are highly hydrophobic, are produced in large amounts and accumulate exclusively in root bark. *L. erythrorhizon* roots have also been utilized as a natural dye to stain clothes by use of a mordant.

*L. erythrorhizon* is a suitable model system to study hydrophobic lipid metabolite secretion for several reasons. First, these cells can be cultured in a cell suspension system capable of producing large amounts of shikonin derivatives (Yazaki et al., 1999). Second, the production of shikonin can be regulated by selection of medium and culture conditions, e. g., shikonin production is strongly induced in M9 medium (Fujita et al., 1981), but is completely inhibited in Linsmaier-Skoog (LS) medium, and light illumination inhibits shikonin production even in M9 medium. In addition, shikonin is exclusively secreted into apoplasts, and is visible as a red pigment under bright-field microscopy and as auto-fluorescence under confocal microscopy.

In this study, we thoroughly analyzed extracellular lipids of cultured *L. erythrorhizon* cells to identify lipid molecules facilitating the secretion of shikonin derivatives, as well as the types of membrane lipids are involved in the secretion of these lipophilic metabolites. We have found that a large proportion of TAG is secreted into extracellular spaces of cultured *L. erythrorhizon* cells, along with several polar lipids and shikonin derivatives. We also found that shikonin-containing oil droplets could be reconstructed *in vitro* with TAG and phospholipids. Taken together, these findings suggest that TAG plays a novel role as a matrix lipid for the secretion of lipophilic specialized metabolites.

## RESULTS

### Behavior of naphthoquinones in living cells

Shikonin is present in *L. erythrorhizon* as ester derivatives with low molecular weight fatty acids (Figure 1A), primarily as acetylshikonin followed by β-hydroxyisovalerylshikonin (Oshikiri et al., 2020). All of these shikonin derivatives are highly hydrophobic and easily crystallized as needles when they are mixed with culture media or water. *L. erythrorhizon* cells producing shikonin derivatives in M9 medium are covered with secreted shikonin derivatives, which appear as numerous red granules in extracellular spaces (Figure 1B). In the presence of light illumination, a condition under which shikonin biosynthesis is strongly suppressed, cells lose their red color but extracellular granules are still present.

**Figure 1.**
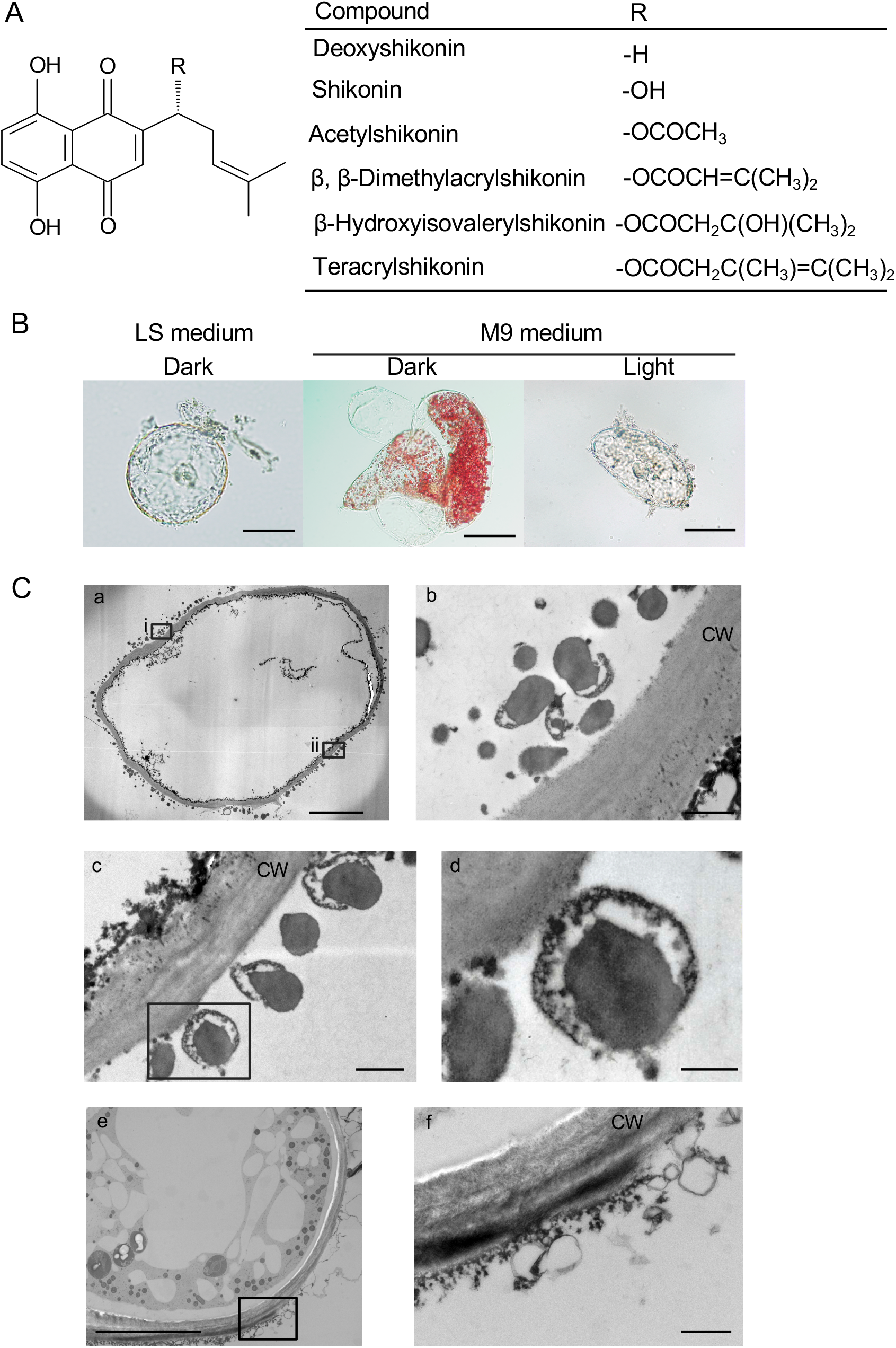
Shikonin-producing cells of *L. erythrorhizon* cultured in M9 medium. (A) Chemical structures of representative shikonin derivatives produced by cultured *L. erythrorhizon* cells. Shikonin has an *R*-configuration at the hydroxyl group on the prenyl chain, with the cultured cells also producing the (*S*)-isomer, called alkannin. (B) Bright-field micrographs of cultured *L. erythrorhizon* cells. LS medium was a negative control, in which shikonin was not produced (left panel). Shikonin derivatives were compartmentalized in the red granules attached to the surface of cells cultured in M9 medium in the dark (middle panel), whereas cells cultured under illumination are incapable of producing shikonin (right panel). Bars = 20 µm. (C) TEM images of shikonin-producing cells of *L. erythrorhizon*. Many characteristic spherical particles were attached to the cell walls. These cultured cells were treated with 5 mM aluminum chloride before fixation and then fixed by standard chemical fixation (a-d) or the HPF/FS method (e, f). Rectangles (i, ii) in (a) depict the enlarged areas in (b) and (c), respectively. Rectangles in (c) and (e) depict the enlarged areas shown in (d) and (f), respectively. Bars = 20 µm (a), 10 µm (e), 1 µm (b, c, f), 500 nm (d). CW, cell wall.

Transmission electron microscopy (TEM) showed characteristic ultrastructures on the walls of *L. erythrorhizon* cells producing shikonin (Figure 1C a). Chemical fixation showed that secreted shikonin derivatives in cultured cells were present as compartments filled with high electron-dense materials (Figure 1C b-d), similar to findings reported in cultured cells (Tsukada et al., 1984). To observe native structures, particularly on membrane structures, cultured *L. erythrorhizon* cells were subjected to high-pressure freezing and freeze substitution (HPF/FS) rather than chemical fixation (Figure 1C e, f and Figure S1). Although HPF/FS showed similar spherical structures of similar size as chemical fixation outside the cells, only the contours of these structures were detected following HPF/FS, because treatment with organic solvents in the latter dissolved and removed the contained materials, including shikonin derivatives. The HPF/FS method also showed two other important differences from chemical fixation: in the HPF/FS method, filamentous structures developed around the cell wall surface to which many spherical membrane structures are attached, and these membrane structures form bunches that fuse with each other to form large inner spaces (Figure S1a-f). These findings suggest that secreted shikonin derivatives are compartmentalized in oil droplets surrounded by membrane structures, thus maintaining the hydrophobic conditions of shikonin derivatives in extracellular spaces.

Even in intracellular spaces, shikonin derivatives are likely sequestered from water-soluble cytosolic components because crystals of shikonin derivatives that are toxic to cells are not observed inside the cells or in apoplasts. Shikonin derivatives have a strong potential to react with water-soluble cellular components (Figure 2A), leading to the gradual decomposition of shikonin (Figure 2B). This decomposition proceeds slowly at room temperature but is accelerated at higher temperature, e.g., 50°C (Figure 2C). Nutrient inorganic cations and amino acids were therefore screened to determine the major components that affect the stability of shikonin derivatives (Figures 2D and 2E). Among cations, Fe was the most probable candidate because shikonin forms a dark purple color in the presence of Fe, which precipitates over time. Although Al and Cu ions also strongly changed the color of shikonin, these cations at the indicated concentrations are not present under physiological conditions or in culture medium, as they are highly toxic to plant cells. Of the 20 amino acids, only cysteine markedly changed the color of shikonin, suggesting that cysteine may be responsible in part for shikonin decomposition upon contact with cell sap (Romero et al., 2014). These findings suggest that Fe and cysteine in the cytosol can react with shikonin derivatives if these naphthoquinone compounds are not sequestered within membrane structures. Shikonin derivatives may coexist with lipid molecules that dissolve these hydrophobic red pigments and are surrounded with membrane lipids in both intracellular and extracellular spaces. We then attempted to analyze lipid molecules synthesized in *L. erythrorhizon* cells.

**Figure 2.**
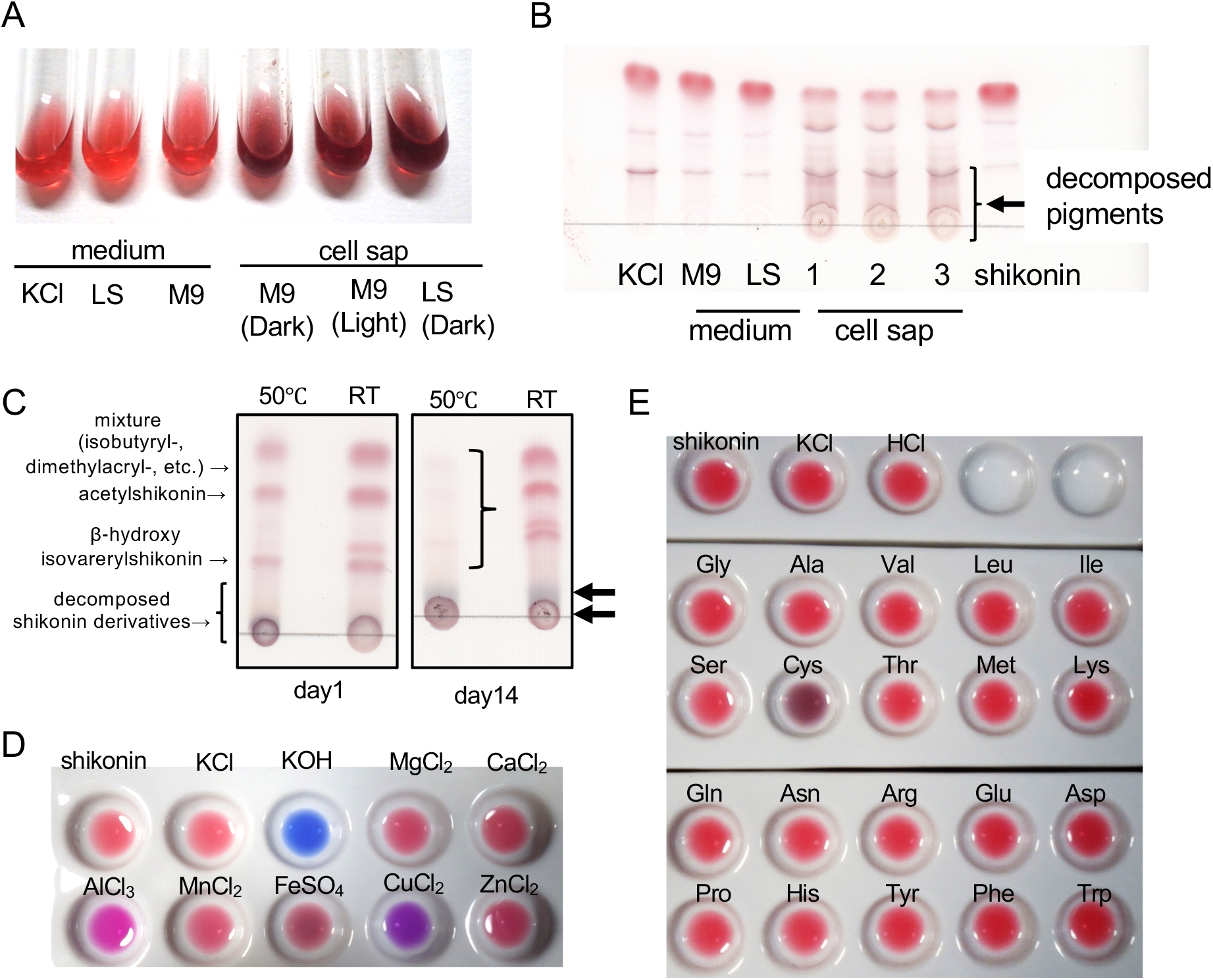
Interactions of cytosolic components with shikonin. (A) Color changes of shikonin samples mixed with fresh culture media or cell sap of *L. erythrorhizon* cells cultured in either medium. KCl was used as a negative control. (B) Thin-layer chromatography (TLC) analysis of shikonin (standard) mixed with fresh culture media or cell sap. Normal phase TLC was developed with 90 : 10 : 1 chloroform : methanol : formic acid. The samples were kept at 25 °C for 2 weeks. The color change reflects the decomposition of shikonin derivatives. (C) TLC analysis (normal phase) of shikonin derivatives coextracted with cell sap (M9 dark culture). The solvent system was as in (B), above. Shikonin derivatives detected are indicated on the left of TLC. The samples were kept at room temperature (RT), or at 50 °C to accelerate the reaction over the time indicated. The parentheses highlight the disappearance of shikonin derivatives and the arrows indicate the generation of decomposed bluish pigments after 14 days at 50 °C. (D) Standard shikonin samples (0.75 mM) were incubated with 25 mM of each inorganic cation. KOH is an alkaline solution used for colorimetric assay of shikonin content. At acidic pH, the color of shikonin was unaffected, with the pH of all solutions of inorganic cations being acidic (pH 3.0–5.7) before mixing with shikonin. (E) Each amino acid was mixed with a standard shikonin sample. KCl and HCl were used as negative controls, with the color of shikonin beings unaffected.

### Lipidomic analysis of extracellular lipid molecules highly produced upon shikonin production of cultured *L. erythrorhizon* cells

To determine the lipid molecules that contribute to shikonin secretion, we performed a comparative lipidomic analysis of *L. erythrorhizon* cells cultured under conditions that enhance (M9 medium, dark) or suppress (M9 medium, light; LS medium, dark) shikonin production. Each batch of cultured cells was divided into three fractions, i.e., medium, cell surface, and cellular fractions as described in the Methods. Total lipids extracted from each fraction with chloroform and methanol were subjected to liquid chromatography-mass spectrometry (LC-MS) analysis. A total of 153 lipid species were detected in culture *L. erythrorhizon* cells, with Figure 3 showing heatmaps of lipid classes detected in these cells. Lipidomic analysis indicated that the production of all lipid classes except TAG was highly activated in shikonin-producing cells cultured in M9 medium in the dark. In particular, the production of phospholipids in the cellular fraction was higher in these shikonin-producing cells than in cells cultured under shikonin non-producing conditions (Figures 3 and S2). Phospholipids are major constituents of cell membranes, including plasma membranes (PM) and organellar membranes. Epidermal cells in shikonin-producing hairy roots of *L. erythrorhizon* contain a highly developed endoplasmic reticulum (ER) network, in which shikonin derivatives are specifically produced (Tatsumi et al., 2016). Analysis of the cellular location of a key shikonin biosynthetic enzyme, *p*-hydroxybenzoate geranyltransferase (PGT), showed that GFP fusion proteins of two paralogues, LePGT1 and LePGT2, had the same distribution pattern as the ER marker GFP-h (Ueda et al., 2010) (Figure S3). This finding suggests that the key step in the biosynthesis of shikonin derivatives occurs in the ER. Importantly, after this reaction, the intermediates acquire hydrophobicity due to the geranyl chain (Yazaki et al., 2002; Ohara et al., 2013). Therefore, the presence of large amounts of phospholipids in shikonin-producing cells is in agreement with the development of the ER, which is responsible for the high production of shikonin. RNA sequencing data indicate that genes encoding glycerol-3-phosphatase acyltransferase (GPAT) are highly expressed in cultured *L. erythrorhizon* cells (Takanashi et al., 2019). These genes, including GPAT1, GPAT4/8 and GPAT9, are involved in the biosynthesis of extracellular lipids, such as cutin, in lysophosphatidic acid (LPA) synthesis from glycerol 3-phosphate and acyl-CoA (Jayawardhane et al., 2018), and in glycerolipid biosynthesis, respectively. These findings also indicate that shikonin-producing cells actively synthesize acylglycerols.

**Figure 3.**
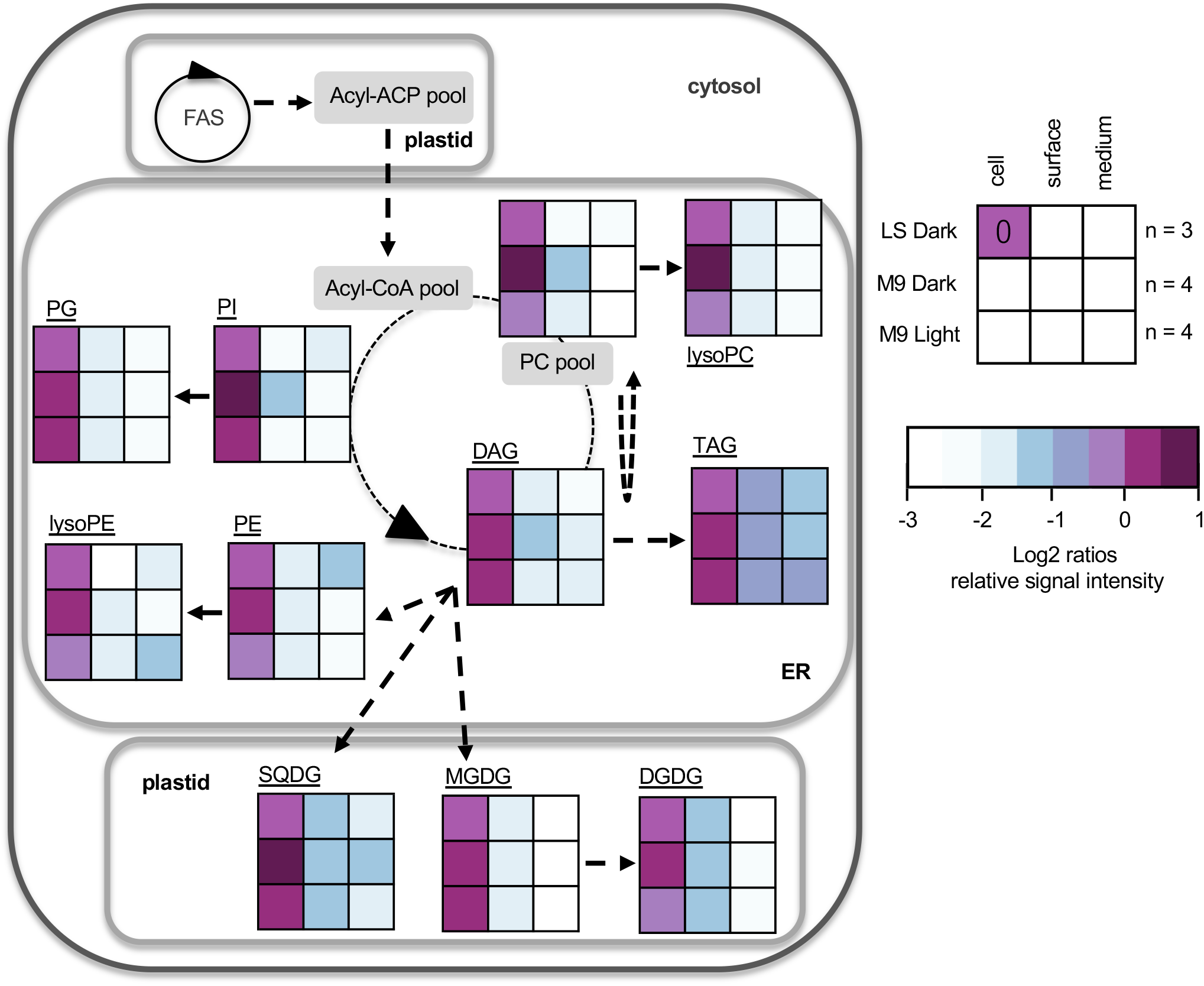
Heat map representation of lipid classes in cultured *L. erythrorhizon* cells along with the glycerolipid metabolic pathway. Average changes in lipid classes are shown by nine boxes (three rows and three columns), which represent three different culture conditions; LS Dark (upper row), M9 Dark (middle row), and M9 Light (lower row); and three fractions: cell, surface, and medium (left-to-right). Heat map colors reveal the average log ratios of fold-changes relative to cell fraction of the LS Dark condition (control). PC, phosphatidylcholine; lysoPC, lysophosphatidylcholine; PE, phosphatidylethanolamine; lysoPE, lysophosphatidylethanolamine; PG, phosphatidylglycerol; PI, phosphatidylinositol; SQDG, sulfoquinovosyldiacylglycerol; MGDG, monogalactosyldiacylglycerol; DGDG, digalactosyldiacylglycerol; TAG, triacylglycerol; DAG, diacylglycerol; Acyl-ACP, acyl-acyl carrier protein; FAS, fatty acid synthase.

Appreciable amounts of TAG were detected outside the cells, both in the medium and cell surface fractions, under all culture conditions, regardless of shikonin production (Figures 3). These results suggest that *L. erythrorhizon* cells secrete large amounts of TAG into extracellular spaces. Generally, TAG accumulates in the cytoplasm as lipid droplets surrounded by a lipid monolayer, a structure conserved in a whole range of organisms, from archaea to mammals, including humans (Murphy, 2012; Ohsaki et al., 2014). This structure is also conserved in higher plants, with the storage of TAG being managed primarily by a universal subcellular organelle called the lipid droplet or oil body (Xu and Shanklin, 2016). Secretion of TAG from cells to extracellular spaces is, therefore, rarely reported in higher plants, but TAG is likely to act as a matrix lipid to solubilize shikonin derivatives in *L. erythrorhizon* cells. Therefore, we further analyzed secreted TAG in detail.

### Cultured *L. erythrorhizon* cells secrete TAG as granular structures into the extracellular space

Field emission scanning electron microscopy (FE-SEM) showed that shikonin-producing cultured cells (M9 Dark) are covered with many small particles, 10–100 nm in diameter, together with larger granules, 1–3 µm in diameter, and filamentous structures (Figure 4A a-c). In contrast, although some large granules are observed on the surfaces of shikonin non-producing cells (LS Dark), few small particles or filamentous structures were present (Figure 4A d-f).

**Figure 4.**
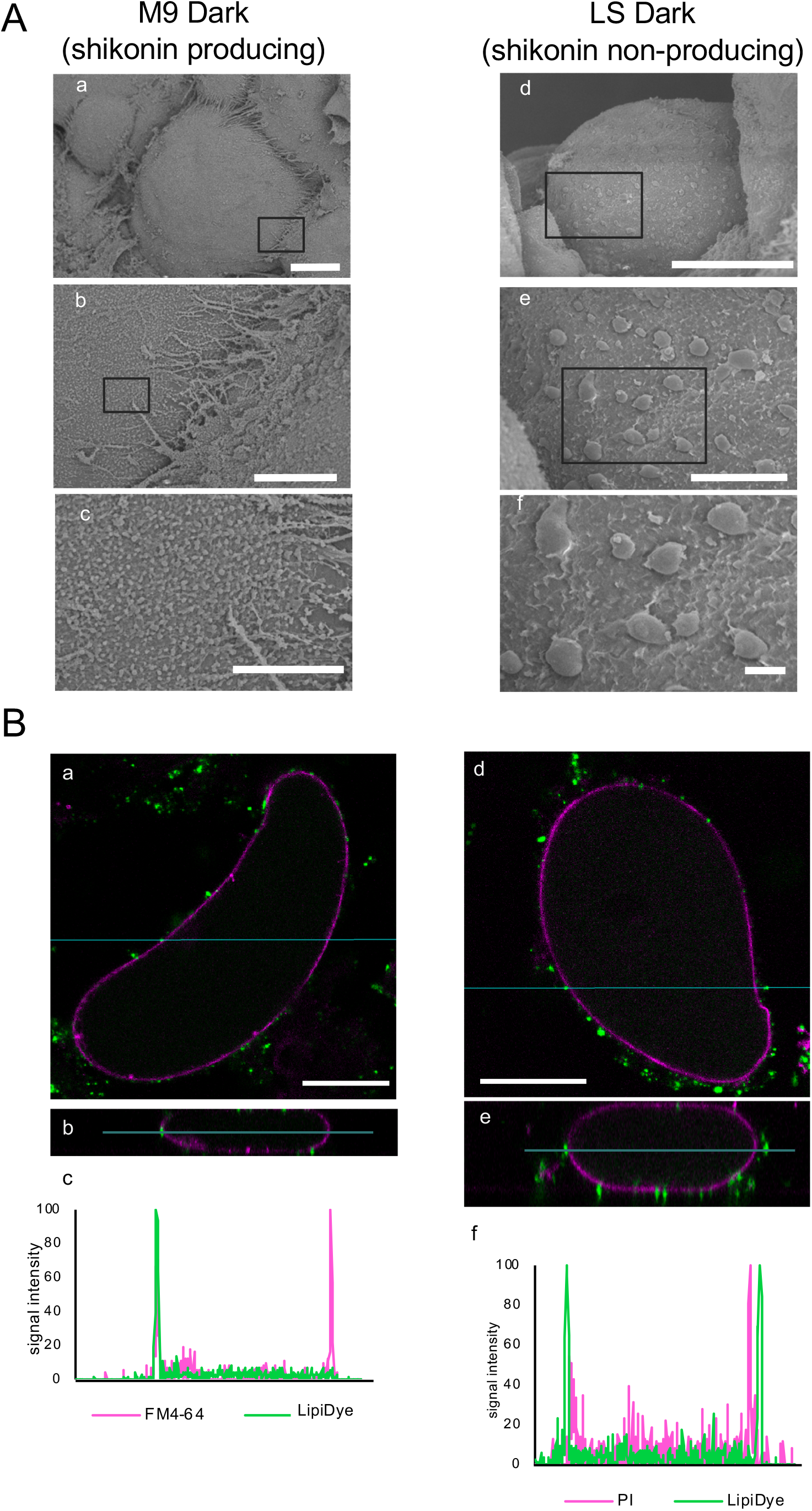
Extracellular particles/granules of cultured *L. erythrorhizon* cells. (A) FE-SEM images of cultured *L. erythrorhizon* cells. (a-c) Shikonin-producing cells cultured in M9 medium in the dark with different enlargements. (d-f) Shikonin-non-producing cells cultured in LS medium in the dark. Panels (b) and (c) and (e) and (f) are enlargements of panels (a) and (d), respectively, with enlarged areas indicated by squares in (a), (b), (d), and (e). Bars = 20 µm (a, d), 5 µm (b, e), 1 µm (c, f). (B) Confocal microscopic images of cultured *L. erythrorhizon* cells stained with LipiDye (pseudocolor green). The PM and cell wall were labeled with FM4-64 (a-c; pseudocolor magenta) and propidium iodide (PI) (d-f; pseudocolor magenta). Z-stack images were captured at 1 μm intervals. Images at xy (a, d) and xz (b, e) views are shown. The blue lines in (a, d) indicate the y axis of (b, e). Signal intensities according to the blue line in (b) and (e) are presented in (c) and (f), respectively. Bars = 50 μm.

To test whether these extracellular granular structures contain secreted lipids, cultured cells were stained with LipiDye, which is specific for lipid droplets, and the cells observed by confocal microscopy (Figure 4B). The location of secreted lipids was analyzed in detail by staining the PM and cell walls with FM4-64 and propidium iodide, respectively. The green fluorescence of LipiDye was detected outside both the FM4-64 and propidium iodide fluorescence, although some LipiDye fluorescence adhered to the PM (Figure 4B a-c) and cell walls (Figure 4B d-f). These results support the above-described findings, that cultured *L. erythrorhizon* cells secrete TAG into the extracellular space, and that this TAG can be visualized as oil droplets.

### Secreted TAG is mainly composed of saturated fatty acids

To quantitatively determine the amount of TAG secreted, TAG was isolated from each fraction of cultured cells by preparative thin-layer chromatography (TLC). TAG appeared between deoxyshikonin and β,β-dimethylacrylshikonin on the TLC plates (Figure S4). Following its isolation from these plates, TAG was analyzed by gas chromatography with a flame ionization detector (GC-FID). About 24-38% of the total amount of TAG produced by the cells was detected in extracellular spaces; i.e., the medium and surface fraction of the cells (Figure 5A). The TAG secretion rate did not differ markedly between shikonin-producing (M9 Dark) and non-producing (M9 Light, LS Dark) cells.

**Figure 5.**
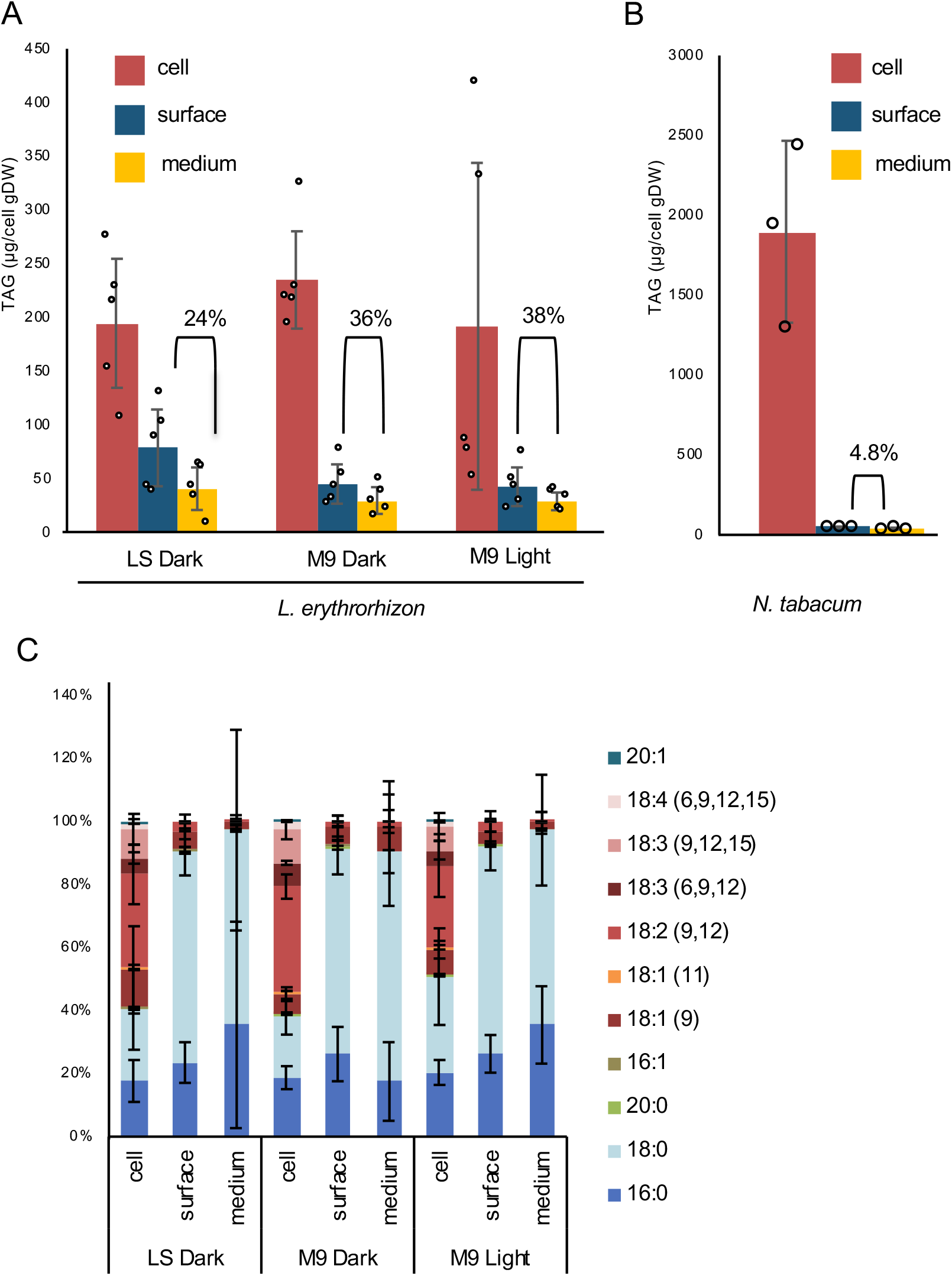
Analysis of TAG molecules secreted by *L. erythrorhizon* cells. (A) Quantitative analysis of TAG in three fractions of *L. erythrorhizon* cells: medium, cell surface, and inside cells. Each point represents the mean of five biological replicates. Open circles represent the individual values of each repeat. Error bars represent the standard deviation (SD). (B) Quantitative analysis of TAG in BY-2 cultured cells. Each point represents the mean of three biological replicates. Open circles represent the individual values of each repeat. Error bars represent the SD. (C) Fatty acid composition of TAG recovered from three fractions of *L. erythrorhizon* cells: medium, cell surface, and inside cells. Each point represents the mean of five biological replicates. Error bars represent the SD.

To investigate whether the secretion of TAG is unique to *L. erythrorhizon* cells or is present in various plant species, TAG amounts were compared in three fractions of cultured *Nicotiana tabacum* cells of strain Blight Yellow-2 (BY-2). GC-FID analysis showed that most of the TAG produced by BY-2 cells accumulated in the cells, with <5% present in the extracellular fractions (i.e., the medium and cell surface) (Figure 5B). These results suggest that the ability to secrete appreciable amounts of TAG into apoplastic spaces is unique to *L. erythrorhizon*.

TAG is an ester compound, composed of a glycerol moiety bound to three fatty acid molecules. The fatty acid composition was therefore compared in secreted TAG molecules and TAG molecules that accumulated inside the cells. TAG in the cellular fraction consisted mostly of unsaturated fatty acids, similar to storage lipids reported in many other plant species. In contrast, however, TAG secreted into the medium and on the cell surface consisted of approximately 90% saturated fatty acids (Figure 5C). The high representation of saturated fatty acids was observed in TAG secreted from both the shikonin-producing and shikonin-non-producing cells, indicating that the fatty acids composing secreted TAG molecules have a unique chemical feature.

### TAG and phospholipids encapsulate shikonin derivatives *in vitro*

Because secretion of the neutral lipid TAG into extracellular spaces implicated that TAG co-existed with shikonin derivatives, the ability of TAG to solubilize shikonin derivatives to form shikonin-containing lipid droplets was assessed *in vitro*. We attempted to prepare lipid droplet-like compartments with TAG and shikonin derivatives in the presence or absence of the phospholipid, phosphatidylcholine (PC). As a control, PC was mixed with triolein, a sort of TAG. The lipid compartment was stained with a neutral lipid-staining dye, BODI PY 493/503, with staining monitored by fluorescence microscopy. This mixture produced small compartments filled with triolein (Figure 6A, B), whereas mixing shikonin with PC did not result in any lipid droplet-like structures (Figure 6C). The mixture of all three components, shikonin, PC, and triolein, yielded shikonin-containing small compartments that could be detected by the auto-fluorescence of shikonin (Figure 6D). To evaluate the effect of the degree of TAG fatty acid saturation on shikonin solubilization, two additional TAG species, trilinolein and tristearin, were tested. Although the three TAG molecules composed of fatty acids of different saturation level showed nearly the same results (Figures 6D-F), the physical properties of these small particles differed slightly, depending on the degree of fatty acid saturation. Lipid particles containing tristearin tended to adhere to the microscope slides and were found to be aggregates of small particles (Figure 6F), suggesting that tristearin-containing lipid particles are more adherent than lipid particles containing unsaturated fatty acids. For further comparison, another fatty acid ester, oleyl linoleate, was employed instead of TAG. However, shikonin was not encapsulated into the particles with oleyl linoleate, but crystallized in the buffer (Figure 6G), suggesting that oleyl linoleate is incapable of solubilizing shikonin, or at least of forming shikonin-containing particles. The mixture of shikonin and triolein without phospholipids yielded a large emulsion of shikonin, which was about 10 times larger than the above-mentioned PC-associated lipid particles (Figure 6H). Moreover, the intensity of shikonin auto-fluorescence of these large droplets was much lower than the intensity observed with the three components, PC, TAG, and shikonin. These results suggest that TAG has the ability to solubilize shikonin and enclose it in an oleophilic compartment. Moreover, membrane lipids, such as PC, are important for encapsulating shikonin in dense particles.

**Figure 6.**
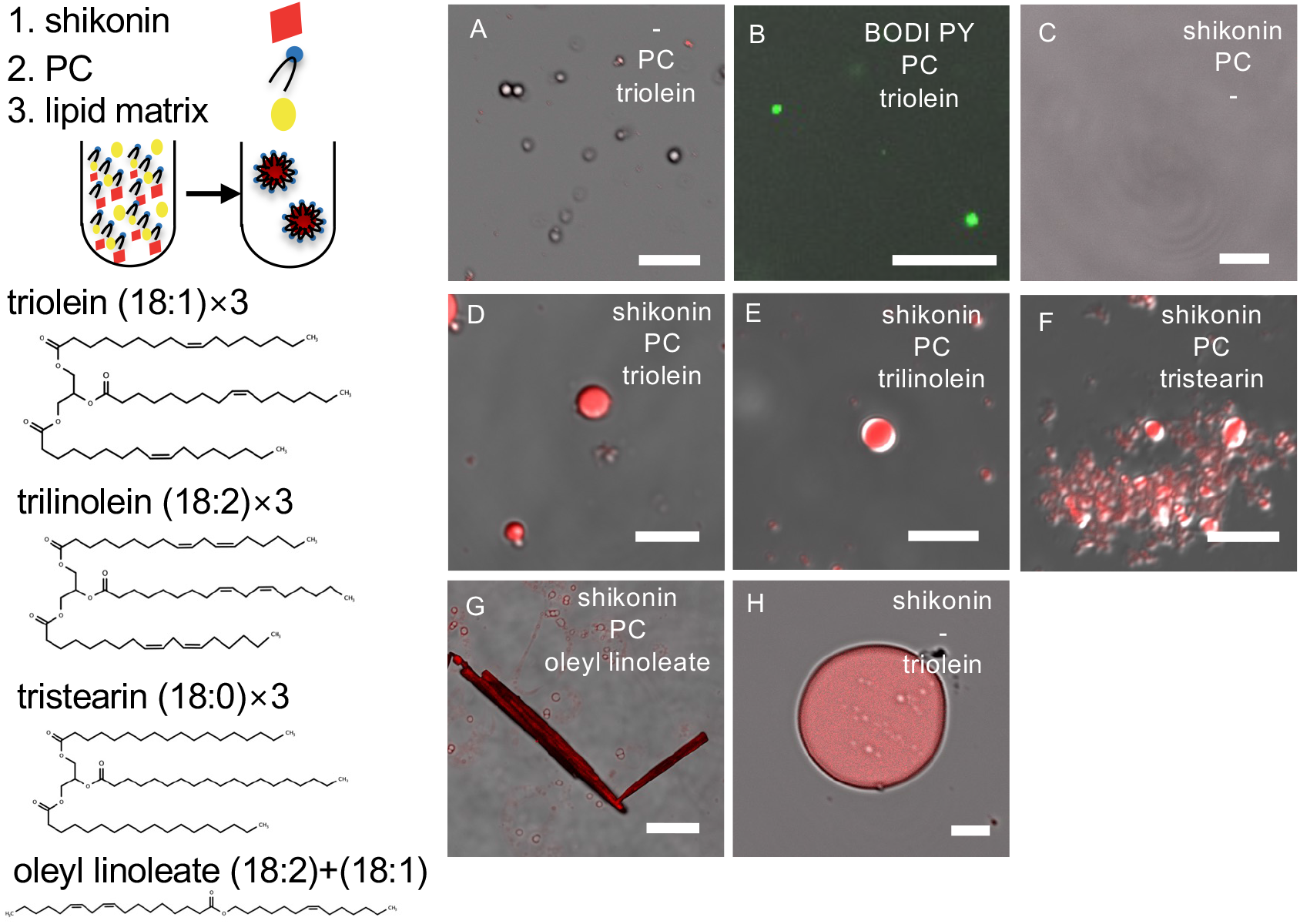
*In vitro* reconstruction of shikonin-containing red droplets with TAG. Lipid particles containing shikonin were constructed by mixing shikonin, TAG, and phospholipid phosphatidylcholine (PC). Three different types of TAG were used; triolein, trilinolein, tristearin. The lipid particles were observed by confocal microscopy. Bars = 20 µm (a, b, c, g, h), 5 µm (d, e, f)

## DISCUSSION

Plants produce a large variety of specialized metabolites and accumulate them in specialized tissues, cells, and organelles (Pichersky and Lewinsohn, 2011; Schenck and Last, 2019). In particular, some of these metabolites, utilized clinically as medicines, show lipophilic properties, such as paclitaxel and vincristine, because of their ability to permeate cell membranes. Lipophilicity is also important for compounds used in perfumes, most of which are volatile organic compounds such as monoterpenes and phenylpropenes, with lower molecular weight compounds being more volatile. Many of these lipophilic specialized metabolites are excreted into apoplastic spaces, where they accumulate. For example, monoterpenes and prenylated phloroglucinols are secreted by secretory cells into the epicuticular cavity in glandular trichomes (Turner et al., 2000). In addition, hydrophobic prenylated flavonoids show similar apoplastic accumulation (Yamamoto et al., 1996). Although much is known about the enzymes responsible for the biosynthesis of these hydrophobic metabolites, little is known to date about the molecular mechanisms underlying the secretion of these lipophilic metabolites by plant cells (Shitan, 2016; Tissier et al., 2017).

The present study used the shikonin production system of cultured *L. erythrorhizon* cells as a model to analyze the secretion of hydrophobic metabolites into extracellular spaces. This study found that TAG is secreted into extracellular spaces, acting as a lipid matrix to solubilize endogenous specialized metabolites, such as shikonin derivatives, in lipophilic compartments surrounded by phospholipids. These results are in agreement with the sequestration of these highly lipophilic naphthoquinone metabolites from cytosol and medium. In this manner, shikonin derivatives do not directly contact cytosolic components like Fe ions or amino acids like cysteine.

In general, TAG is recognized as a storage lipid that accumulates in the cytosol as ‘oil bodies’, intracellular organelles often observed in sink organs (Shimada et al., 2018). These organelles provide a source of energy and precursors of primary metabolites during seed germination, thereafter playing a pivotal role in the early growth of seedlings. TAG is also responsible for the production of ATP, which is essential for stomatal opening in guard cells (McLachlan et al., 2016). The present study also suggests a new role of TAG, acting as a lipid carrier to facilitate the secretion of endogenous hydrophobic specialized metabolites from the cytosol to extracellular spaces. Shikonin biosynthesis occurs on ER membranes; i.e., after the geranylation of *p*-hydroxybenzoic acid, the intermediate is hydroxylated by P450 and finally esterified with an acyltransferase (Figure S3) (Yazaki et al., 2002; Oshikiri et al., 2020; Wang et al., 2019). Utilization of TAG as a matrix lipid is probably advantageous, because the biosynthesis of TAG also takes place on ER membrane, thus allowing both shikonin, and TAG to interact with each other to form shikonin-containing oil droplets.

Lipidomic and GC-FID analyses showed that *L. erythrorhizon* cells could secrete appreciable amounts of TAG (ca. 30% of total TAG produced) to extracellular spaces irrespective of shikonin production (Figure3 and 5A), suggesting that these cultured cells constitutively secrete TAG. Because only epidermal cells in intact roots of this plant species can produce shikonin, the cultured cells are dedifferentiated maintaining the ability to produce and secrete shikonin derivatives. In contrast to shikonin biosynthesis, which is sensitive to illumination and ammonium ion, TAG synthesis and secretion by these cells are insensitive to both (Yazaki, 2017). FE-SEM showed that the number of extracellular particles was much higher in shikonin-producing cells than in shikonin non-producing cells (Figure 4A). Large amounts of membrane lipids are required to fill the entire surface area of lipophilic particles, in agreement with the increase in phospholipids following the induction of shikonin production in M9 medium (Figure 3).

In addition to cultured cells, hairy roots of *L. erythrorhizon* can secrete TAG (Figure S5). Few studies to date have assessed TAG secretion in higher plants. For example, the surface of bayberry (*Myrica pensylvanica*) fruits is covered with a thick lipid layer, consisting primarily of TAG (Simpson et al., 2016). More generally, epidermal cells of terrestrial plants have the ability to secrete lipid compounds, such as the polymers cutin and wax, which protect plant bodies from dryness. Shikonin derivatives are also secreted exclusively by epidermal cells, perhaps by a pathway also responsible, at least in part, for the secretion of TAG. The TAG synthesis pathway may have evolved as an adaptation of cutin synthesis (Simpson and Ohlrogge, 2016). TAG may also be secreted by particular cells or tissues, other than those of *L. erythrorhizon*, from which lipophilic metabolites are secreted. A charophytic alga *Klebsormidium flaccidum* also secretes TAG, suggesting the need for studies of the evolutionary aspects of TAG secretion (Kondo et al., 2016).

Detailed analysis of its fatty acid composition revealed that TAG secreted by *L. erythrorhizon* is mainly composed of saturated fatty acids (Figure 5C). This finding is in agreement with the secreted TAG of bayberry fruits, which are also mainly composed of saturated fatty acids (Simpson and Ohlrogge, 2016). Aliphatic components of polymers in cuticular wax are synthesized from saturated, very long-chain fatty acids (Kunst and Samuels, 2003). The reason for the difference in secreted and stored TAG species is unclear, but their biosynthetic pathways likely differ, similar to findings in bayberry fruits (Simpson et al., 2016). TAG composed of saturated fatty acids may have biological advantages. Lipid particles containing tristearin are stickier than those containing triolein and trilinolein (Figure 6D-F). Many shikonin-containing particles adhere to the surface of in shikonin-producing cultured cells (Figure 1B middle panel). In intact *L. erythrorhizon* plants, shikonin derivatives accumulate at the boundary between root tissues and soil. By sticking to the root surface, secreted shikonin-containing droplets tightly cover root tissues acting as a chemical barrier. In addition, shikonin derivatives were reported to have antimicrobial properties (Brigham et al., 1999).

In recent years, plant extracellular vesicles have been actively studied. Extracellular vesicles are broadly defined as lipid-enveloped particles containing proteins, RNAs, and other metabolites (Cai et al., 2018; Pegtel and Gould, 2019; Liu et al., 2020). These vesicles, which can be classified by origin, function, and or size (diameter, 30-10,000 nm), are thought to participate in various plant defense mechanisms (Rutter and Innes, 2017; Regente et al., 2017). Extracellular shikonin-containing particles observed by TEM and FE-SEM in this study are 10–100 nm in diameter, with bright light and fluorescence microscopy showing that they grow larger on the cell surface, to 1–3 µm, over time. Because shikonin derivatives are thought to be defense compounds, these shikonin-containing particles may be similar to extracellular vesicles. However, due to their high hydrophobicity, shikonin-containing particles may be similar to lipid-monolayer droplets, growing larger via fusion with each other in apoplastic spaces as observed under bright microscopy (Figure 1B middle panel). The present findings provide novel examples of lipid-containing extracellular vesicles in plants.

In conclusion, the present study showed that shikonin derivatives, as a model of extracellular hydrophobic specialized metabolites, are secreted with TAG from cells as droplets that are encapsulated by membrane lipids. These results provide perspectives for the role for TAG, which functions as carrier lipid for endogenous specialized metabolites to transport them to extracellular spaces. The rapid development of new analytical technologies has increased studies on the biochemistry and molecular biology of shikonin in *L. erythrorhizon* and related species (Wang et al., 2019; Ueoka et al., 2020; Oshikiri et al., 2020; Izuishi et al., 2020; Yamamoto et al., 2020; Song et al., 2021), including the sequences of their genomes (Tang et al., 2020; Auber et al., 2020). Further studies are needed to evaluate the molecular mechanisms by which TAG and shikonin are secreted at the plasma membrane, and the types of proteins involved in the secretion process.

## Limitations of the Study

This study had several limitations. First, we were unable to analyze direct interactions between shikonin derivatives and TAG *in vivo*. To analyze the effects of lipid desaturation on shikonin secretion, we attempted to generate transgenic *L. erythrorhizon* hairy roots, in which endogenous fatty acid desaturase 2 (LeFAD2) and stearoyl-ACP desaturase 1 (LeSAD1) were overexpressed or knocked down. However, we could not obtain any cell lines overexpressing these enzymes under the control of an estrogen inducible promoter (Zuo et al., 2000), and RNAi hairy roots did not grow on growth medium.

## Resource Availability

### Lead Contact

Requests for further information and for resources should be directed to and will be fulfilled by the Lead Contact, Kazufumi Yazaki.

## Materials Availability

Materials generated in this study are available from the Lead Contact with a completed Materials Transfer Agreement.

## Data and Code Availability

The datasets supporting the current study have not been deposited in a public repository because they are parts of further investigations. However, they are available from the corresponding author upon request.

## METHODS

### Plant materials and growth condition

Cultured cells (strain T-TOM) of *L. erythrorhizon* were maintained in Linsmaier-Skoog’s (LS) medium, containing 3% sucrose, 10^−6^ M potassium indole acetate (Nacalai Tesque, Kyoto, Japan) and 10^−5^ M kinetin (Sigma, St. Louis, MO) at 25 °C, 80 rpm in darkness, with cells subcultured at 2-week intervals. To induce shikonin production, these cultured cells were transferred to M9 medium containing 3% sucrose (Fujita et al., 1981), 10^−6^ M potassium indole acetate and 10^−5^ M kinetin, and cultured in the dark or under illumination with the same agitation conditions.

Hairy roots of *L. erythrorhizon* were induced as described (Tatsumi *et al*., 2020), and hairy root cultures were maintained in 1/2 Murashige-Skoog’s medium with 3% sucrose at 25 °C, 80 rpm in the dark, with hairy roots subcultured at 2-week intervals. For shikonin production, these hairy roots were cultured in M9 medium without plant hormones for two weeks. Tobacco BY-2 (*Nicotiana tabacum* L. cv. Bright Yellow-2) cells were cultured in modified LS medium at 28 °C, 100 rpm in the dark, with cells subcultured at weekly intervals (Nagata et al., 1992).

### Interaction of shikonin with water-soluble components of cell sap

Cells cultured for 2 weeks in M9 medium in the dark or light, or in LS medium were subjected to a freeze-thaw process (frozen at -20℃ and thawed at room temperature for 30 min) thrice to destroy membrane structures. Each sample was homogenized with a spatula in a glass vessel, which was centrifuged at 500 rpm for 1 min to obtain the cell sap as supernatant. A 300 µL aliquot of standard shikonin solution (1 mM in 1-propanol) was mixed either with 100 µL cell sap and allowed to stand for 2 weeks in the dark at room temperature. As a negative control, an aqueous solution of 100 mM KCl was used instead of cell sap. Each medium was also employed for comparison.

Interactions of shikonin with cell sap was analyzed by normal phase TLC (silica gel 60 F254, Merck Millipore, Darmstadt, Germany). In addition, a 20 µL aliquot of each sample was spotted onto a TLC plate, which was developed with a 90 : 10 : 1 mixture of chloroform : methanol : formic acid.

The interactions of shikonin (standard) with metal ions were evaluated by mixing 1 mM shikonin solution in 1-propanol with aqueous solutions of 100 mM KCl, MgCl_2_, CaCl_2_, AlCl_3_, MnCl_2_, FeSO_4_, CuCl_2_, and ZnCl_2_ at a 3 : 1 (v/v) ratio. Because the red color of shikonin turns blue under alkaline conditions, the aqueous solution of each inorganic salt was confirmed to be at acidic pH, from pH 3.03 for FeSO_4_ to pH 5.73 for MgCl_2_, indicating that any color change was not due to alkaline pH. As a control for blue color, 1 mM shikonin was mixed 3 : 1 (v/v) with 2.5% KOH.

Similarly, the interactions between shikonin and amino acids were evaluated by mixing 1 mM shikonin solution in 1-propanol with aqueous solutions of 100 mM of each amino acid at a 3 : 1 (v/v) ration. Because tyrosine, tryptophan, aspartic acid, and glutamic acid had to be dissolved in diluted HCl, as they were poorly soluble in water, diluted HCl was used as a negative control, as was KCl.

The effect of alcohol on the stability of shikonin derivatives in the extract was evaluated by the direct extraction of pigments from crude dried roots of *L. erythrorhizon*. Briefly, the dried roots were soaked in 95% ethanol, which was maintained at room temperature or at 50 °C. Extracts were spotted onto TLC plates after 1 and 14 days, and the plates were developed with a solvent system composed of n-hexane : acetone : formic acid (80 : 20 : 1).

### Lipid extraction

Lipids of plant cells and tissues (cultured cells and hairy roots of *L. erythrorhizon* and BY-2 cells) were extracted into three parts, *i*.*e*., medium, cell surface, and the rest of the cell fraction. The liquid cultures were filtered through Miracloth (Merck Millipore) to separate the culture medium (30 ml) from the cells or root tissues. The cultured medium was partitioned with 15 ml of 2 : 1 (v/v) chloroform : methanol to yield organic phase (medium fraction). The harvested wet cells/tissues were rinsed with 15 ml of 2 :1 (v/v) chloroform-methanol and 30 ml distilled water by prompt everting of the glass vessel to recover the cell surface lipids (surface fraction). The remaining cells/tissues were completely dried and the lipids extracted with 2 ml of 2 :1 (v/v) chloroform-methanol to yield the cell fraction. Each fraction was evaporated under nitrogen stream before chromatographic analyses.

### LC-MS analysis

Before LC-MS analysis, lipid extracts were roughly separated into polar and non-polar lipids by thin-layer chromatography (TLC) using silica plates (TLC silica gel 60, Merck Millipore) developed with chloroform because the high amount of shikonin derivatives hampered the chemical analysis of TAG and polar lipids by LC-MS. Lipid samples, except for shikonin derivatives that could be recognized by their red color, were recovered from TLC plates and extracted with chloroform or methanol from the silica gel. The lipids were subjected to LC-q-TOF-MS (Waters, Boston, MA) analysis with an Acquity UPLC HSS T3 column (Waters), as described (Okazaki et al., 2013; Okazaki and Saito, 2018). Lipidomic analysis was performed using the data set recorded in the positive ion mode. Levels of lipid species were normalized relative to the intensity of the internal standard PC (20:0). To compare the amount of each lipid class among samples, the level of each lipid class, which is the sum of individual lipid species belonging to the class, was standardized by using the mean of cell fraction in LS Dark cultured cells.

### GC-FID analysis

To quantify TAG, TAG was purified by preparative TLC developed with 6 : 4 (v/v) n-hexane : diethyl ether. Following derivatization to fatty acid methyl esters using a fatty acid methylation kit (Nacalai Tesque), TAG was quantified by capillary gas chromatography GC-2014 (Shimadzu, Kyoto, Japan) with a J&W DB-23 capillary column (GL Science, Tokyo, Japan) as described (Kajikawa et al., 2015; 2016), with heptadecanoic acid (C17:0) used as the internal standard.

### DNA cloning

cDNA of cultured *L. erythrorhizon* cells was synthesized (Tatsumi et al., 2020) using KOD Plus Neo DNA polymerase (Toyobo, Osaka, Japan). An open reading frame (ORF) eliminating stop codon of the *LePGT2* gene was amplified, and the fragment was subcloned into pENTR/D-TOPO (Invitrogen, Carlsbad, CA) by in-fusion reaction (Clontech, Mountain View, CA) using a linearized vector. To obtain the plant expression vector for mGFP fusion, the cDNA fragment was cloned into the destination vector pGWB405m (Nakagawa et al., 2007; Segami et al., 2014) using LR clonase II (Invitrogen). Primers used for vector construction are listed in Table S1.

### LePGT1 and LePGT2 subcellular localization

Plant vectors expressing LePGT1-mGFP (Tatsumi et al., 2020) and LePGT2-mGFP, as well as GFP-h (ER marker) (Ueda et al., 2010), were introduced into *Agrobacterium tumefaciens* (strain LBA4404) by the freeze-thaw transformation method. Each was transiently expressed in *N. benthamiana* leaves by agroinfiltration using transformed agrobacteria at OD_600_ 0.1. Two days after infection, GFP fluorescence was detected.

### Staining with fluorescent dyes

Stock solutions of LipiDye (1 mM; Funakoshi, Tokyo, Japan) and FM4-64 (40 mM; Invitrogen) were prepared in dimethyl sulfoxide, and a stock solution of propidium iodide (1 mg/mL; Wako, Osaka, Japan) was prepared in water. Cultured *L. erythrorhizon* cells were incubated with 2 µM (final concentration) LipiDye for 2 hours, and then with 80 µM FM4-64 or 6 µg/mL propidium iodide (final concentrations for both). To remove excess amounts of fluorescent dyes, the cells were gently washed with phosphate buffered saline. The cultured cells were monitored immediately after washing.

### Light microscopic analysis

Light microscopic pictures of cultured *L. erythrorhizon* cells were captured by a Axioscope 2 (Zeiss, Oberkochen, Germany).

### Confocal microscope analysis

LePGT-GFP of *N. benthamiana* leaves was monitored by excitation at 488 nm with a 20 mW diode laser and emission at 500–540 nm. Images of *L. erythrorhizon* cultured cell and lipid-containing particles were observed using a confocal laser scanning microscope FV3000 (Olympus, Tokyo, Japan) with a 60 × 0.75 numerical aperture water immersion objective. LipiDye was monitored by excitation at 405 nm with a 50 mW diode laser and emission at 521–530 nm. FM4-64 and propidium iodide were monitored by excitation at 561 nm with a 40 mW diode laser and emission at 570–670 nm. The autofluorescence of shikonin was monitored by excitation at 561 nm with a 20 mW diode laser and emission at 570–670 nm. BODI PY 493/503 (Molecular Probes, Eugene, OR) was monitored by excitation at 488 nm with a 20 mW diode laser and emission at 500–540 nm. Image acquisition and analysis were performed using FV31S-SW software and ImageJ software (http://rsb.info.nih.gov/ij). Signal intensity was determined by the Plot Profile tool of ImageJ.

### Transmission electron microscopy

Cultured cells were treated with 5 mM aluminum chloride for 3 h before chemical fixation of shikonin derivatives, as described (Tatsumi et al., 2016). High-pressure freezing and the freeze substitution method (HPF/FS) were performed as follows. Cells cultured in M9 or LS medium containing 3% sucrose were added to flat specimen carriers and frozen using a high-pressure freezing machine (Leica EM PACT, Leica Microsystems, Wetzlar, Germany). The frozen samples were transferred to 2% osmium tetroxide in anhydrous acetone at -80 °C and incubated at -80 °C for 6 days (120 h). These samples were warmed gradually from -80 to -30 °C over 5 h, warmed again from -30 to 4 °C over 3.4 h, and held at 4 °C for 2 h (Cryo Porter CS-80CP, Scinics Corporation, Tokyo, Japan). Subsequently, the samples were washed with acetone, and embedded in Epon812 resin (TAAB, Aldermaston, England). Ultrathin sections (60-80 nm) were cut with a diamond knife on an ultramicrotome (Leica EM UC7, Leica Microsystems) and placed on formvar-coated copper grids. The ultrathin sections were stained with 4% uranyl acetate followed by lead citrate solution and observed with a JEM-1400 (JEOL Ltd., Tokyo, Japan) transmission electron microscope at 80 kV. Some sections were not stained with lead citrate to prevent excess staining.

### Scanning electron microscopy

Cultured cells were fixed with 2% glutaraldehyde, 50 mM sodium cacodylate buffer (pH 7.4) for 2 h at room temperature and then for 4 d at 4 ºC. After post-fixation with 1% osmium tetroxide, 50 mM sodium cacodylate buffer, the samples were dehydrated in a graded methanol series (12.5, 25, 50, 75, 90, and 100%). The samples were infiltrated with isopentyl acetate, dried with a critical point dryer (CPD-030, Bal-tec, Balzers, Liechtenstein), and coated with osmium tetroxide in an osmium coater (HPC-1SW, Vacuum Device, Mito, Japan). The cells were subsequently monitored with a field emission scanning electron microscope (SU8220, Hitachi High-Tech, Tokyo, Japan) at 3 kV.

### Construction of lipid emulsion

Lipid-containing particles were constructed as described (Arisawa et al., 2016), modified to allow the addition of shikonin derivatives to the particles. To construct shikonin-containing particles, PC (Wako) in methanol, either TAG molecule in 2 : 1 (v/v) chloroform : methanol, and standard shikonin in chloroform were mixed in a molar ratio of 1 : 20 : 105. After drying under a nitrogen stream, the residue was resuspended in 150 mM NaCl, 50 mM Tris-HCl (pH 7.5), 1 mM EDTA, 1 mM dithiothreitol, and 50 mM phenylmethylsulfonyl fluoride solution and sonicated for 10 sec three times (duty cycle 70%, output control 3) with a sonicator (Sonifier 250, Branson, Danbury, CT). The resulting lipid-containing particles were harvested by ultracentrifugation at 100,000 g for 15 min at 37 °C. Four types of neutral lipids were tested: triolein (C18:1×3), trilinolein (C18:2×3), tristearin (C18:0×3) and oleyl linoleate (Wako).

## ACKNOWLEDGMENTS

The authors thank Dr. Hirobumi Yamamoto (Toyo University) for providing cultured *L. erythrorhizon* cells and Dr. Noboru Onishi (Kirin Holdings Company Limited) for providing *L. erythrorhizon* axenic shoots, which were used to generate hairy roots. The authors also thank Dr. Nam-Hai Chua (Rockefeller University) and Dr. Takashi Aoyama (Kyoto University) for providing the pTA7002 and pER8GW vectors. The authors also thank Dr. Takahiro Hamada (Okayama University of Science), Dr. Haruko Ueda, and Dr. Ikuko Hara-Nishimura (Konan University) for providing the pBI121_GFP-h vector; and Dr. Shoji Segami (National Institute for Basic Biology) and Dr. Masayoshi Maeshima (Chubu University) for providing the pGWB405m vector.

The authors also thank Ms. Mayumi Wakazaki (RIKEN CSRS) and Mr. Kenta Kaminade (Kyoto University) for technical assistance with electron microscopy and plant cell cultures, respectively.

This work was funded by the Japan Society for the Promotion of Science (JSPS) KAKENHI to K.Y. (JP19H05638), a JSPS Research Fellowship for Young Scientists DC2 to K.T. (201811502), and the New Energy and Industrial Technology Development Organization (NEDO) to K.Y. (16100890-0).

## AUTHOR CONTRIBUTIONS

K. Tatsumi, T.I., A.S., and K.Y. designed the research; K. Tatsumi., T.I., N.I., Y.O., Y.H., M.K., and M.S. performed the experiments. K. Tatsumi, T.I., H.F., K. Toyooka, analyzed the data. H.O., K. Saito, and I.I. contributed new technical and analytical methods. K. Shimomura provided *L. erythrorhizon* axenic shoots for hairy root generation. K. Tatsumi, and K.Y. wrote the paper with critical input from K. Toyooka and M.S.

## Supplemental Figures

**Figure S1**. Transmission electron micrographs of cultured *L. erythrorhizon* cells fixed with the HPF/FS method. Enlarged parts of a cell in (a) are indicated with white squares (b - d). Panels (e, f) are of different cells from (a), showing frequently seen bunches of characteristic extracellular structures. Panels (g) and (h) highlight small electron-dense particles inside the cell walls, in which cultured cells were treated with 5 mM aluminum chloride before fixation to fix shikonin derivatives. CW, cell wall. Scale bars are: a, 2 μm; b, d, e, 500 nm; c, g, h, 200 nm; f, 1 μm.

**Figure S2**. Relative signal intensity of each lipid class. The values of fraction in LS Dark cells (shikoinn-non-producing) were set to 1 for each lipid class. Black dots represent individual values of each repeat (n = 3; LS Dark, n =4; M9 Dark and M9 Light). PC, phosphatidylcholine; lysoPC, lysophosphatidylcholine; PE, phosphatidylethanolamine; lysoPE, lysophosphatidylethanolamine; PG, phosphatidylglycerol; PI, phosphatidylinositol; SQDG, sulfoquinovosyldiacylglycerol; MGDG, monogalactosyldiacylglycerol; DGDG, digalactosyldiacylglycerol.

**Figure S3**. Subcellular localization of LePGT expressed in *N. benthamiana* epidermal cells. Confocal microscopic images of LePGT1-mGFP (A), LePGT2-mGFP (B), and GFP-h (C) transiently expressed in epidermal cells of *N. benthamiana* leaves by agroinfiltration. Bars = 10 µm

**Figure S4**. Normal phase thin-layer chromatography (TLC) analysis of lipid fraction from *L. erythrorhizon* cells cultured in M9. The TLC was developed with n-hexane: diethyl ether (6 : 4, v/v). (Left) Without staining, with shikonin derivatives appearing as red spots; (right) following staining with I_2_ vapor after separation, showing TAG spots. Shikonin derivatives identified in standard specimens are also indicated on the right.

**Figure S5**. Normal phase thin-layer chromatography (TLC) analysis of lipids extracted from hairy roots of *L. erythrorhizon*. The TLC was developed with n-hexane: diethyl ether (7: 3, v/v). (Left) Without staining; (right) after staining with 80% (v/v) aqueous acetone containing 0.01% (w/v) primuline. Yellow arrowheads indicate TAG spots.

**Table S1**. Primers used for LePGT2 gene construction.

