## Supplementary Figures for "Excretion of triacylglycerol as a matrix lipid facilitating apoplastic accumulation of a lipophilic metabolite shikonin"

### Supplemental Figures

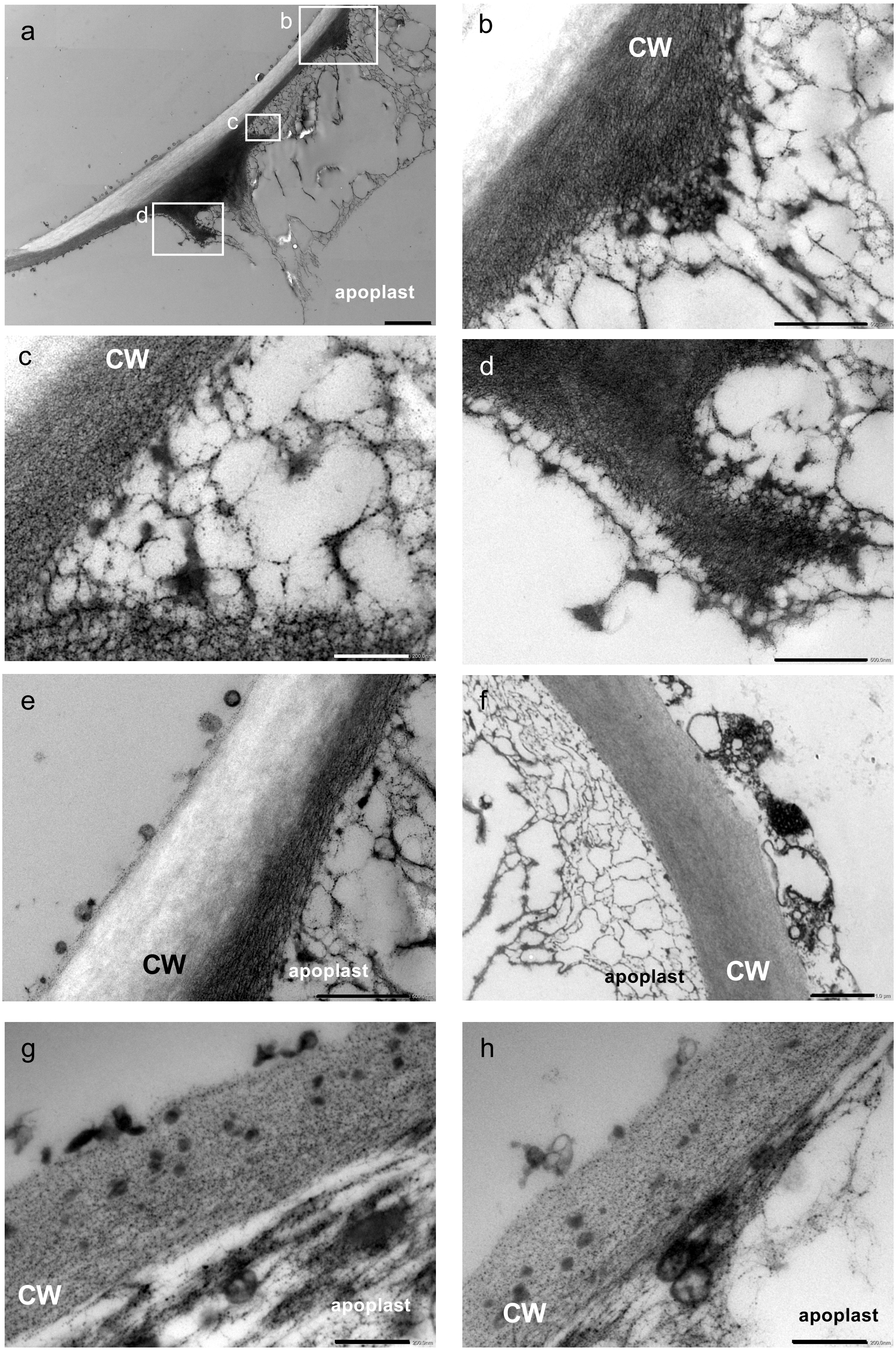


Supplemental Figure. 1

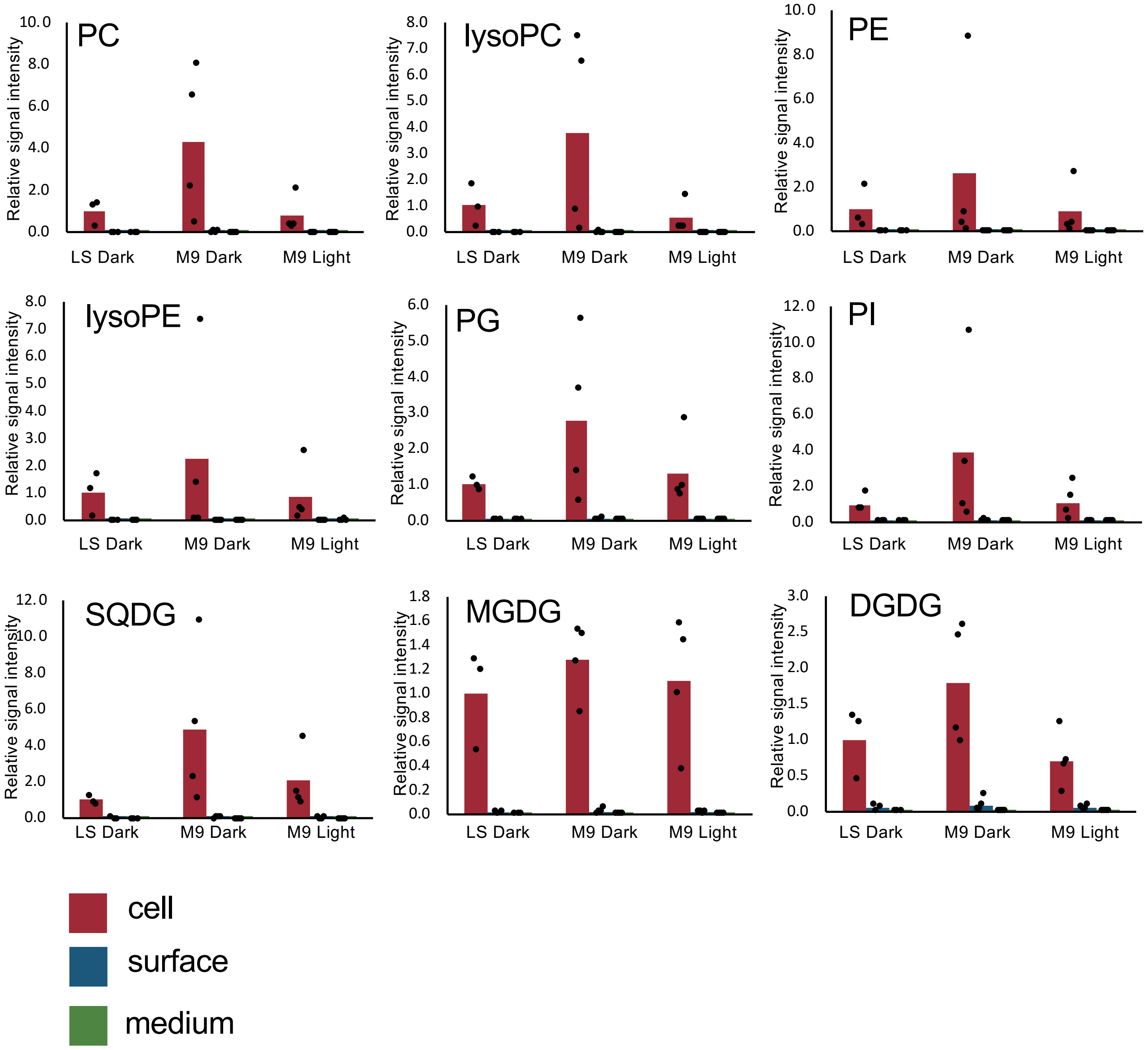


Supplemental Figure. 2

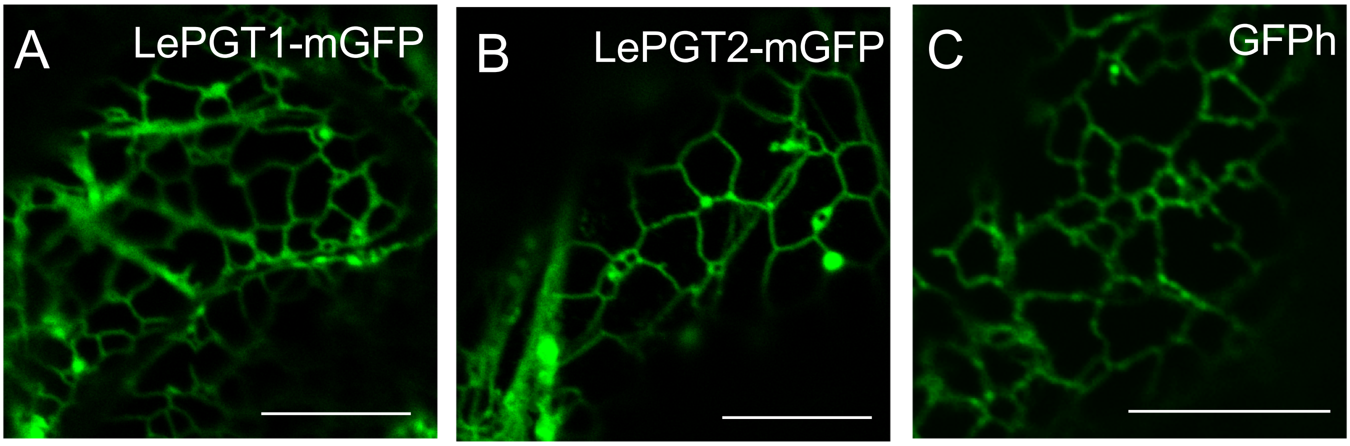


Supplemental Figure. 3

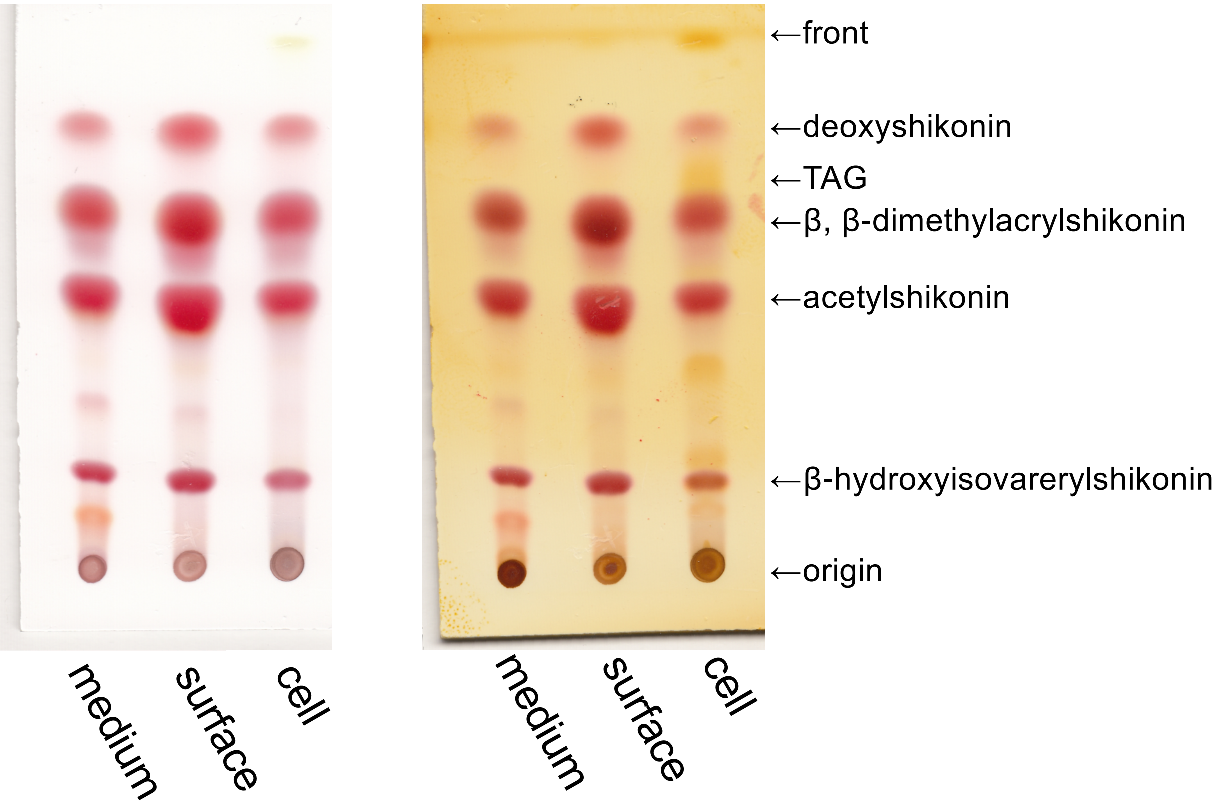


Supplemental Figure. 4

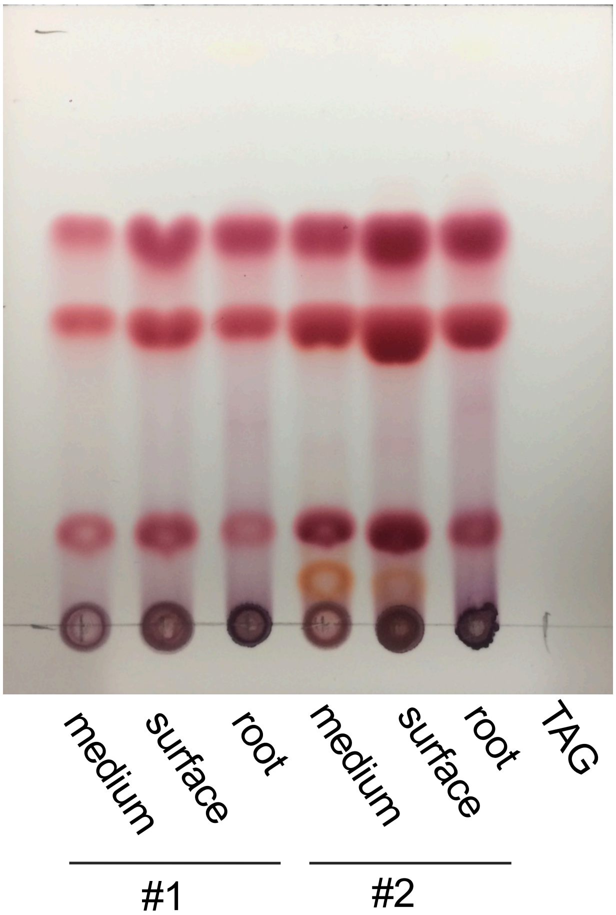


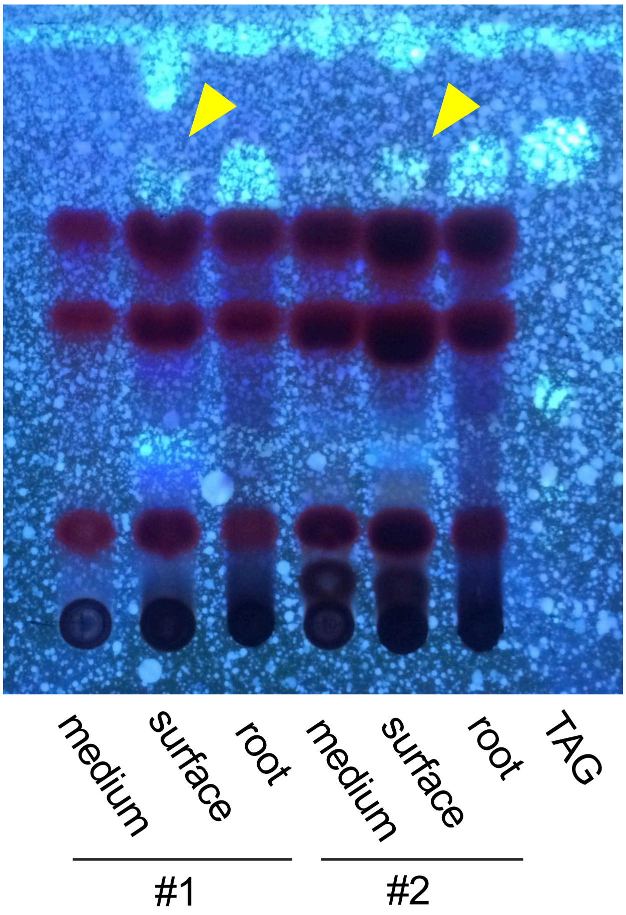


Supplemental Figure. 5

Supplemental Table 1. Primer list

| purpose for use | gene/vector | sequence |
|---|---|---|
| cloning of LePGT2 using In-fusion reaction | LePGT2 | Fw: GCAGGCTCCGCGGCCGCCACCATGA GTTC CAAACAAACACAGCTAAAGAA |
| | | Rv: AGCTGGGTCGGCGCGCCCAGGAAAC AATC TCCCAACTAAGATGC |
| linearizing pENTR/D-TOPO | pENTR/D-TOPO | Fw: CGCGCCGACCCAGCTTTCTTGTACAA AGTT |
| | | Rv: GGCCGCGGAGCCTGCTTTTTTGTACA AAGT |
